# Method for maintaining translocated wild roses under laboratory conditions for controlled gall induction by *Diplolepis rosae* and *D. mayri*

**DOI:** 10.64898/2026.05.19.726150

**Authors:** Zoltán László, Avar-Lehel Dénes, Sarah Melissa Witiak, Eszter Péterfi, Dorina Podar

## Abstract

1. Plant-gall wasp systems provide unique models for studying multitrophic interactions and unique developmental processes, yet standardized laboratory protocols for maintaining wild rose hosts (*Rosa* spp.) and sustaining gall inducers (*Diplolepis* spp.) are lacking.
2. We developed and tested a method for growing and maintaining translocated individuals of *Rosa canina, R. rubiginosa, R. spinosissima, R. gallica, R. tomentosa*, and *R. pendulina* under laboratory conditions over three consecutive years (2023–2026) to provide a continuous source of host plant material for experimental gall induction by *D. rosae* and *D. mayri*.
3. The protocol integrates soil and substrate composition, photoperiod and humidity regimes, pruning, dormancy management, and controlled exposure to gall-inducing wasps. More than 75% of rose individuals survived the full 3-year period, with consistent annual gall induction achieved in several host species.
4. This work represents the first reproducible laboratory method for long-term maintenance of wild rose hosts and controlled gall induction by *Diplolepis* species. The framework also provides a broadly applicable approach for maintaining perennial woody hosts and experimentally manipulating specialized plant–insect interactions under laboratory conditions, thereby supporting ecological, physiological, and evolutionary research across comparable systems.

## 1. INTRODUCTION

Gall induction by cynipid wasps (*Diplolepis* spp.) on wild roses is a conspicuous and ecologically significant plant-insect interaction in temperate ecosystems (Stone & Schönrogge, 2003). A range of gall morphologies are produced when female wasps oviposit in very young leaf tissues and developing buds (Stone, 1998). Gall structures provide food and shelter for their inducers and associated communities of parasitoids, inquilines, and fungi (Shorthouse & Rohfritsch, 1992). While gall midges (Cecidomyidae) dominate plant gall diversity, in the northern hemisphere there is also a high diversity of gall wasp (Cynipoidea) inducers on trees, shrubs and herbs (Dorchin et al., 2019; Hayward & Stone, 2005). Compared to herb cynipid galls, those on trees and shrubs exhibit well-defined tissues with diverse and complex morphologies (Stone, 1998; Stone & Schönrogge, 2003). Among them, *Diplolepis rosae* (the “bedeguar gall wasp”) and *D. mayri* are widespread in Europe, inducing large mossy or spiny spherical and multilocular galls on wild *Rosa* species (László et al., 2024).

Although natural populations of wild roses and gall inducers can be monitored in situ, experimental studies are often constrained by seasonality, limited access, and the unpredictability of gall induction in the field. For instance, while numerous studies explore the transcriptomes of fully developed galls (Takeda, 2019; Martinson, 2022), only a few explore the early stages of gall induction on woody plants (Schultz, 2019) because of the difficulty to precisely identify early induction plant tissue sites, or collect consistent replicates in the field. Long-term maintenance of host plants under controlled conditions could provide year-round experimental access to not only monitor gall induction, but also manipulate the galling process to test ecological, evolutionary or molecular hypotheses (László et al., 2024; Murakami, 2021). Not all woody species are suited to laboratory conditions, but roses are one example of galling susceptible woody plants that can be maintained in a small area and multiplied via clonal propagation to produce replicates.

For the commercial rose market, numerous methods for rose cultivation exist (Bredmose, 1992; Davarzani, 2023). These protocols generally focus on efficient flower production from continuously flowering cultivars, unlike wild roses which usually flower only once a year.

However, in our case, rose gall induction in the lab requires the reliable production of vegetative buds and young leaves. Here we present a standard laboratory method for growing and maintaining multiple wild rose species over consecutive years while ensuring reliable gall production.

We grew translocated *Rosa canina, R. rubiginosa, R. spinosissima, R. arvensis, R. gallica, R. tomentosa*, and *R. pendulina* under controlled laboratory conditions from 2023 to 2026. We describe plant establishment and maintenance, followed by controlled introduction of gall wasps to induce galls. We evaluate plant survival, growth, and gall induction success, and provide recommendations for future studies.

## 2. MATERIALS AND METHODS

### 2.1. Source and translocation of plant material

Wild individuals of the seven different *Rosa* species were collected as young shoots with root systems from natural populations in Transylvania (Romania) during spring 2023, 2024 and 2025 (Table 1). Plants were transported with native soil and transplanted into ∼0.5L containers filled with a semi-standardized soil mixture (30% native soil, 50% potting soil for indoor plants (AGRO CS, Romania), 15-18% river sand (particle size 0.06-2 mm), 2-5% fertilizer).

**TABLE 1.** List of wild rose collecting sites from Romania. N: number of collected individuals.

| Species | Year | Location, county code | N |
| --- | --- | --- | --- |
| *R. canina* | 2024 | Ciumbrud, AB | 2 |
|  | 2023, 2024 | Luna de Jos, CJ | 12 |
|  | 2023, 2024, 2025 | Cluj-Napoca, CJ | 11 |
| *R. rubiginosa* | 2023 | Ciumbrud, AB | 1 |
|  | 2023, 2024 | Luna de Jos, CJ | 7 |
| *R. spinosissima* | 2024 | Moldovenești, CJ | 2 |
|  | 2024 | Cheile-Turzii, CJ | 2 |
| *R. gallica* | 2023, 2024 | Ciumbrud, AB | 10 |
|  | 2023, 2024 | Luna de Jos, CJ | 64 |
| *R. arvensis* | 2024 | Sfânta Elena, CS | 4 |
|  | 2024 | Dubova, MH | 5 |
| *R. tomentosa* | 2025 | Sfânta Elena, CS | 3 |
| *R. pendulina* | 2025 | Bologa, CJ | 3 |

### 2.2. Growth and maintenance conditions

The growth parameters were selected to simulate in situ late spring and early summer conditions, when roses are actively producing new vegetative buds that provide sites for gall induction.

Environmental conditions were maintained year-round at a constant 21±2°C, 60-70% air relative humidity, and a constant 16hr:8hr (light:dark) photoperiod. Full spectrum LED lights (40W Cosmorrow, Secret Jardin, Agomoon SRL, Belgium, www.secretjardin.com) provided 120 - 300 µmol/m^2^/s with wavelengths between 440 and 750 nm. Permanent air ventilation was maintained using ∼40W air towers to prevent fungal growth. Pots were placed in shallow trays to ensure a permanent 3-5 mm water level to maintain soil humidity. The soil mix in the pots was renewed each winter to sustain plant vigor. These conditions supported year-round vegetative growth.

Nutrient supplementation consisted of different fertilizers: for the first two years a fast-release fertilizer (AGRO liquid fertilizer for roses, NPK 4-6-7, AGRO CS Hungary Kft. (www.agrocs.hu) was applied three times per year, while in the last year a slow-release fertilizer (Dr. Soil Solid Fertilizer for Roses, NPK 2-2-4, Dr. Soil GmbH, www.doctorsoil.com) was applied twice per year.

During maintenance time, infestations by mites, aphids and scale insects occurred. To control aphids and mites, plants were washed 1-2 times per week under a strong spray of water. Scale insects were removed with a brush under running water or a cotton swab dipped in 70% ethanol. Where gall inducers and galls were not present, we used chemical control against aphids, mites and fungi when necessary (Triple Action Ikebana, Quimica de Mungia S.A. Spain, www.quimunsa.com).

Plants were pruned twice per year: in spring, two months before oviposition, to stimulate ramification and bud development, and in autumn (September-October) after the collection of mature galls, to stimulate new shoot growth. Additional pruning was applied as needed to fast-growing individuals to maintain plant sizes compatible with growth chamber limitations. Plants with newly emerged radicular-bud-derived stems were divided at the base in spring 2024 to produce genetic replicates.

### 2.3. Gall induction by *Diplolepis* spp

In late spring (May-June) of each year (2023-2026), plants were exposed to adult *D. rosae* and *D. mayri* collected from local field populations. Each potted rose was enclosed within a tulle mesh (25□× □10□× □10□cm) (Figure 1). A single newly emerged female wasp was released onto the rose plant beneath the tulle mesh. Dead wasps were removed 4-7 days after the exposure. Gall development was monitored over the subsequent 8–12 weeks (László et al., 2025).

**FIGURE 1.**
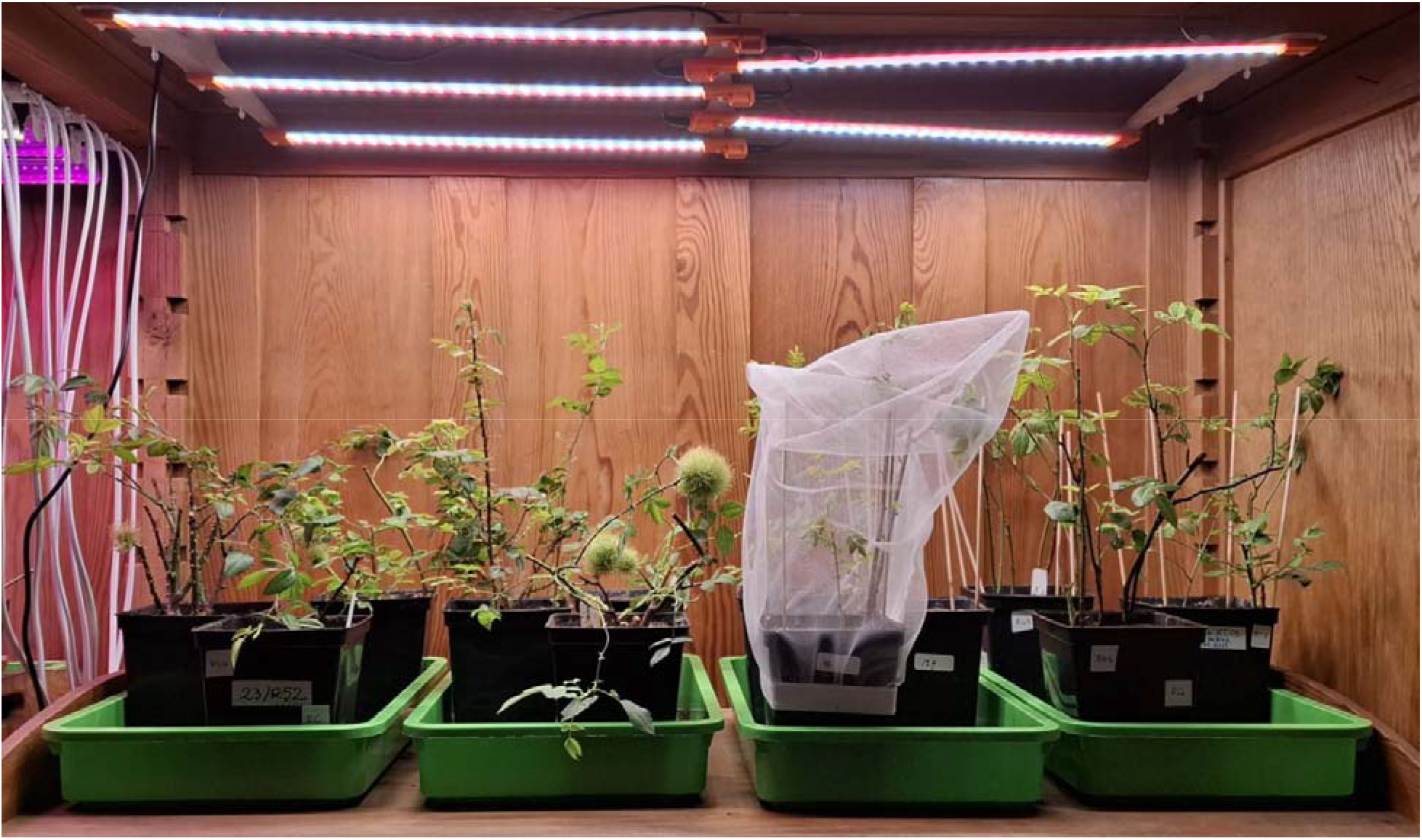
Potted wild roses under LED lights with enclosed tulle meshes and growing rose galls.

### 2.4. Monitoring and data collection

For each rose species the number of galls per wasp species were recorded (Table 2). Gall induction success was defined as ≥1 gall formed on a given plant during the season.

**TABLE 2.** Number of wild rose individuals successfully galled (2023-2025).

|  | *D. rosae* |  |  | *D. mayri* |  |  |
| --- | --- | --- | --- | --- | --- | --- |
|  | 2023 | 2024 | 2025 | 2023 | 2024 | 2025 |
| *R. canina* | 2 | 9 | 11 | 1 | 5 | 1 |
| *R. rubiginosa* | 2 | 5 | 1 | 1 | 0 | 0 |
| *R. gallica* | 0 | 0 | 1 | 0 | 0 | 0 |

### 2.5. Statistical analysis

Survival was analysed using a binomial generalized linear model with logit link, fitted in R (R Core Team, 2026) using the stats package (glm). The response was specified as counts of alive and dead individuals. Fixed effects included time (continuous) and *Rosa* taxon. Initial maximum-likelihood models showed quasi-complete separation for some taxa, producing unstable estimates; therefore, a bias-reduced GLM was additionally fitted using the brglm2 package (Kosmidis, 2021) to obtain more reliable parameter estimates. Model terms were assessed using Wald z-tests. Predicted survival probabilities were generated on the response scale across the observed time range using the ggplot2 package (Wickham, 2016) for visualisation. Nagelkerke’s pseudo-R^2^ was used to quantify the proportion of variance explained by the fixed effects in the binomial logistic regression model (Nagelkerke, 1991).

## 3. RESULTS

### 3.1. Plant survival and growth

Wild rose survival differed among the studied *Rosa* species over the course of the observation period (Figure 2A). Overall survival between the beginning of the study in 2023 and the final assessment in 2026 ranged from 50% in *Rosa arvensis* and *R. pendulina* to 100% in *R. rubiginosa* and *R. gallica*. Intermediate survival rates were observed for *R. canina* (around 85%), *R. spinosissima* (80%), and *R. tomentosa* (75%). When examining survival across individual observation years (Figure 2B), several species maintained consistently high survival, particularly *R. rubiginosa* and *R. gallica*, which remained at 100% throughout the monitoring period. In contrast, *R. canina* showed a gradual decline in survival from the first to the third year, while *R. spinosissima* also decreased between the first and second year. *R. tomentosa* exhibited comparatively a lower survival value (75%) during the year it was observed, whereas *R. pendulina* and *R. arvensis* showed the lowest survival (50%). Overall, there were clear interspecific differences in both cumulative survival and temporal survival dynamics among the studied *Rosa* species. Indoor growth conditions maintained the plants in a vegetative state throughout the three-year period, and no flowering was observed.

**FIGURE 2.**
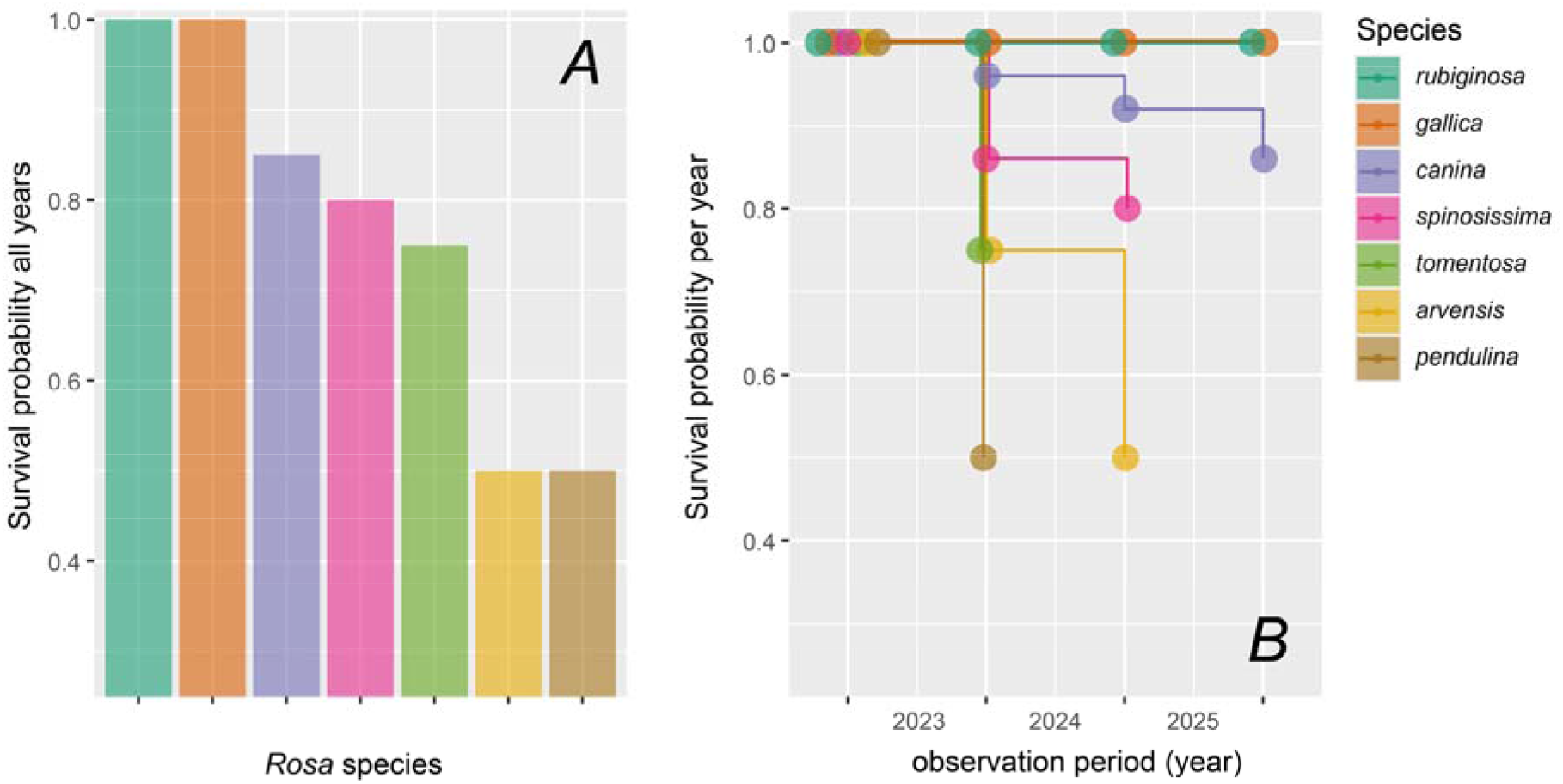
Survival of the studied *Rosa* species during the observation period: A) overall percentage survival of individuals from the beginning of the study in 2023 to the final assessment in 2026 for each species; B) annual survival percentages for each species across one-, two-, and three-year observation periods, highlighting temporal changes in survival among species.

### 3.2. Gall induction success

The number of galls varied among both *Diplolepis* species and their host *Rosa* species over the three-year observation period (Figure 3). While *D. mayri* produced the majority of galls in 2023 and 2024, its abundance declined markedly by 2025, when most recorded galls were associated with *D. rosae* (Figure 3A). *Rosa canina* hosted an increasing number of galls throughout the study. In contrast, galls on *R. rubiginosa* were common in 2023 but decreased substantially in subsequent years, whereas *R. gallica* hosted only a small number of galls and only in the latest year of the study (Figure 3B). The data indicate a general increase in *D. rosae* galls on *R. canina* over time, while galls associated with *D. mayri* and *R. rubiginosa* declined markedly (Figure 3C). These patterns demonstrate clear shifts in gall abundance among both *Diplolepis* species and their *Rosa* hosts during the study period.

**FIGURE 3.**
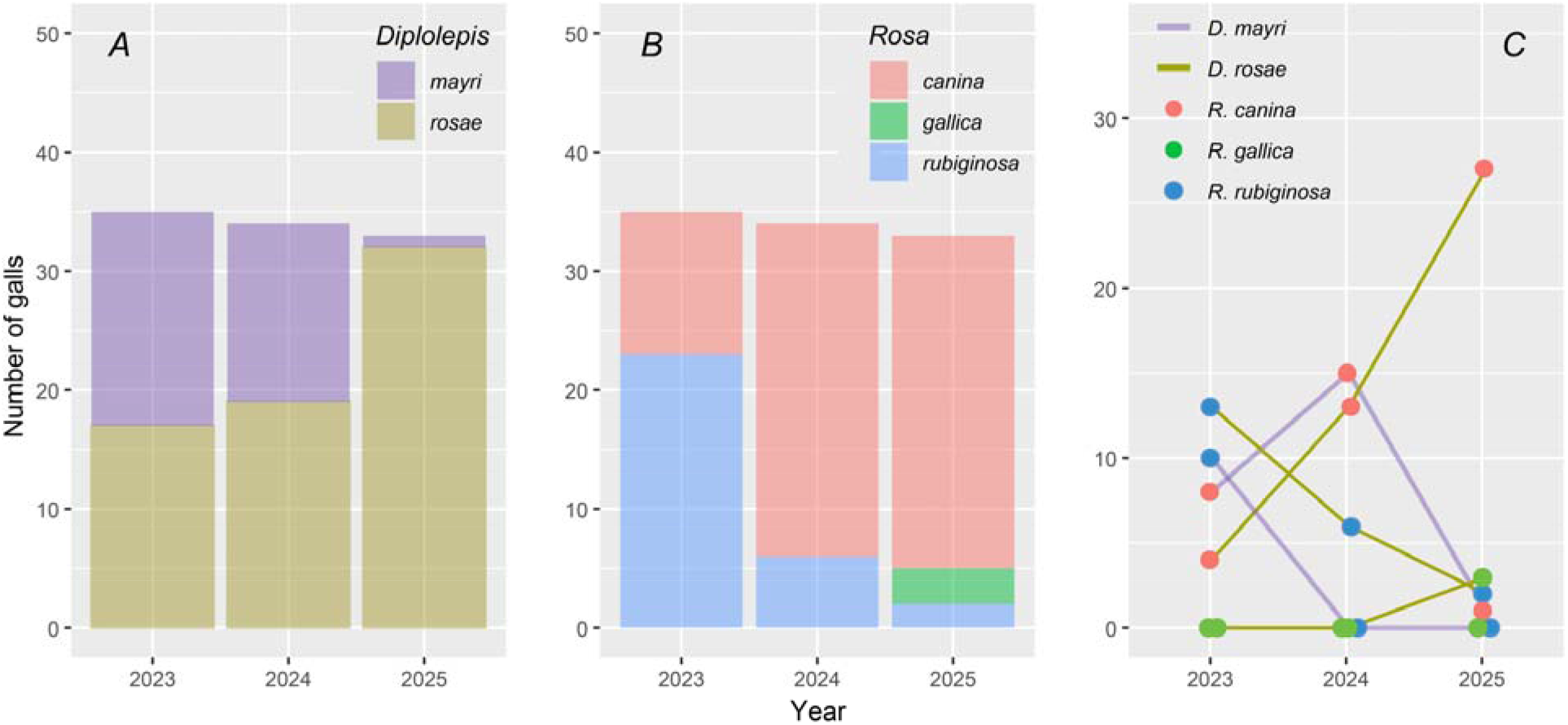
Gall abundance associated with *Diplolepis* species and their *Rosa* hosts from 2023 to 2025: A) the total number of galls produced by the two gall-inducing wasps, *Diplolepis mayri* and *D. rosae*, for each study year; B) the number of galls recorded on each rose species (*Rosa canina, R. gallica*, and *R. rubiginosa*); C) temporal changes in gall abundance across the three years, showing the dynamics of galls associated with both *Diplolepis* species and their respective *Rosa* hosts.

### 3.3. Multi-year consistency

Survival probability with time and taxon (*Rosa* spp.) as predictors, indicated quasi-complete separation (notably for *R. gallica* and *R. rubiginosa*) based on bias-reduced estimates. There was a temporal trend in rose survival (β = −1.06 ± 0.37, z = −2.85, p = 0.004). Survival also varied among taxa, compared to the survival of *R. pendulina*, which had the lowest survival rate: *R. canina* (β = 3.19 ± 1.04, p = 0.002), *R. gallica* (β = 6.32 ± 1.71, p < 0.001), and *R. rubiginosa* (β = 4.63 ± 1.75, p < 0.001) all showed significantly higher survival. The species *R. spinosissima* showed a weaker, non-significant increase in survival (β = 1.79 ± 1.03, p = 0.081). Similarly, *R. arvensis* (β = 0.64 ± 1.00, p = 0.522) and *R. tomentosa* (β = 0.65 ± 1.18, p = 0.583) which showed also no significant differences from *R. pendulina*. Model fit was good (χ^2^ = 39.24, df = 7, p < 0.001), and predicted survival curves were consistently separated across time, indicating stable interspecific differences and strong multi-year consistency for some of the species (Nagelkerke’s R^2^ = 0.83, Figure 4).

**FIGURE 4.**
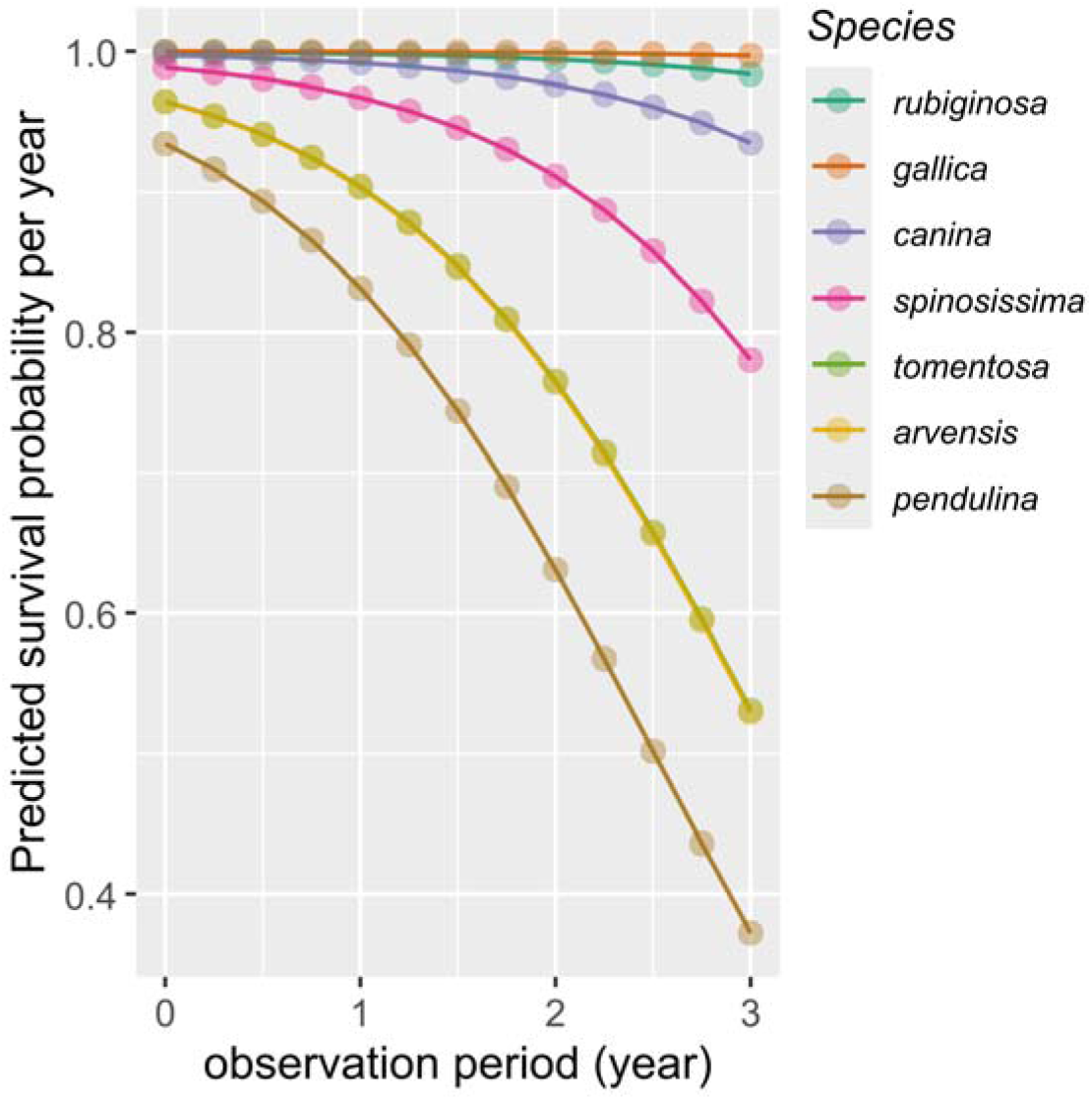
Predicted survival probability over time for different *Rosa* taxa based on a bias-reduced binomial generalized linear model. Lines show model-predicted probabilities across the observed time range for each species.

## 4. DISCUSSION

Our study provides the first reproducible method for long-term laboratory maintenance of wild rose hosts for gall induction. The protocol successfully sustained multiple *Rosa* species over three consecutive years, demonstrating the feasibility of year-round experimental availability of host plants. Key elements contributing to success included: controlled conditions to maintain the vegetative state; optimization of soil mixtures, minimal fertilizer input and regular pruning to ensure consistent vegetative growth and to maintain manageable plant sizes.

This made it possible to study the controlled exposure of wild roses to gall-inducing wasps in tulle mesh enclosures. Other attempts to induce galls on commercial wild *Rosa* species were unsuccessful, possibly because the purchased plants hybridized with cultivated varieties (pers. comm.). This highlights the importance of careful identification of species that have not hybridized with cultivated roses.

Survival probability varied across years, indicating that the suitability of laboratory-maintained rose hosts is not fully constant over time. This likely reflects changes in plant condition, physiological state, or other environmental factors affecting host quality, and suggests that temporal variation is an important component of the system. In this context, active and consistent plant care is likely required to maintain stable host quality. At the same time, clear and relatively large differences among *Rosa* species were observed, with *R. canina, R. gallica*, and *R. rubiginosa* showing the highest survival probabilities. These species therefore appear to provide more suitable developmental conditions for *Diplolepis* wasps. Overall, host identity remains the main driver of gall induction success, while temporal variation should be considered when interpreting outcomes, consistent with previous findings that host quality strongly affects oviposition and larval performance in plant-insect interactions (Stone et al., 2002).

All of the wild rose species collected are able to be grown in the greenhouse. While the survival rate differed between species, *R. canina* and *R. rubiginosa* proved to be persistent and able to support galls in the lab. The ability to induce two different types of galls upon 3 different wild rose species provides a useful system to test various ecological and developmental hypotheses by comparing these different interactions. The wild rose species that did not produce galls will be useful in the future to test gall resistance or for insect choice studies.

The *R. canina* – *D. rosae* interaction is particularly robust in the lab under the described conditions which may be of help in studying broad ecological questions. For example, *R. canina* habitat is expected to decline by ∼ 30% with climate change (Arslan, 2020). Presumably galling interactions will also be affected by climate change. The development of a laboratory-friendly galling system for this species will allow validation of climate change scenarios.

Beyond *Rosa*–*Diplolepis* systems, the methodology developed here has potential for broader application across plant–insect interaction research. The framework provides an approach for maintaining perennial woody hosts and experimentally manipulating specialist herbivore interactions under laboratory conditions, and may be adaptable to other gall-inducing, endophagous, or tightly host-associated insect systems used in ecological, physiological, and evolutionary research. In particular, controlled vegetative maintenance of wild host plants, targeted gall induction under enclosed conditions, and multi-year stability assessment may be applicable to systems involving specialist herbivores dependent on specific host phenologies.

Similar protocols also may be adapted for cynipid wasps on oak (*Quercus* spp.), a well-studied system where host identity and phenology effects galling success (Stone & Schönrogge, 2003), or for leaf-mining insects that require consistent host tissue availability for larval development (Lopez-Vaamonde et al., 2020). The challenge of sourcing and maintaining non-hybridized wild host plants, highlighted here for *Rosa*, is equally relevant in any study system where cultivated or hybrid variants may differ in defence chemistry or developmental cues. More broadly, the integration of long-term greenhouse maintenance with controlled oviposition arenas may serve as a template for laboratory domestication of other ecologically important but experimentally intractable plant-insect systems, supporting research in chemical ecology, plant defence, and community-level interactions such as hyperparasitism and inquilinism.

This approach opens opportunities for experimental studies on gall formation, plant-insect defence physiology, insect developmental plasticity, and multitrophic interactions in *Rosa-Diplolepis* systems. It may also serve applied research, such as testing the ecological roles of gall communities under climate change scenarios.

## 5. CONCLUSION

We developed a standardized method for growing and maintaining wild *Rosa* species under laboratory conditions over a 3-year period, ensuring consistent gall induction by *D. rosae* and *D. mayri*. The method provides a foundation for ecological, physiological, and evolutionary studies on rose-gall interactions, comparable in utility to established laboratory systems in insect-plant research.

## ACKNOWLEDGMENTS

We acknowledge the support of the greenhouse staff at Babeş-Bolyai University for their technical assistance during this study. In particular, we thank Zoltán Balázs. We thank Attila Mátis for identifying wild rose species. We are grateful for student assistance from Ágnes Balázs, Anna Ercse, Mátyás Illyés, and Róbert Veres. Sarah M. Witiak was supported by a Fulbright Scholar Award 2025-2026.

## AUTHOR CONTRIBUTIONS

Zoltán László designed the study, coordinated plant maintenance, and wrote the manuscript. Sarah M. Witiak and Lehel Dénes-Avar contributed to plant maintenance, data collection and analysis, and wrote the manuscript. Eszter Péterfi assisted with plant maintenance. Dorina Podar contributed to conceptualization, method development, and manuscript review.

## DATA AVAILABILITY STATEMENT

The data that support the findings of this study are available on request from the corresponding author. The data are not publicly available due to privacy or ethical restrictions.

